# Exercise Intensity Modulates the Validity of Non-Linear Heart Rate Time Series Analysis Window Length: Implications for DFAa1 Monitoring

**DOI:** 10.64898/2026.02.02.703217

**Authors:** J. De Maeseneer, A. Olieslagers, T. Gronwald, T. de Beukelaar

## Abstract

**Purpose:** Detrended fluctuation analysis alpha-1 (DFAa1) has emerged as a promising non-invasive biomarker for exercise intensity assessment. However, the standard 2-min analysis window lacks temporal resolution necessary for real-time training applications. This study systematically investigated the validity of shortened DFAa1 windows (30s and 1min) versus the 2-min reference across different intensities.

**Methods:** Physically active males completed three continuous cycling protocols: low-intensity training at the first lactate threshold (LOW, n=19), moderate-intensity training at the second lactate threshold (MOD, n=19), and a 30-min self-paced time trial (TT_30_, n=18). DFAa1 was calculated using 30-s, 1-min, and 2-min moving windows, advancing in 1s increments. Validity was assessed using intraclass correlation coefficients (ICC), Bland-Altman analysis, and standard error of measurement (SEM).

**Results:** During LOW, both shortened windows showed poor agreement with the 2-min reference (30s: ICC=0.02, mean bias of –0.05; 1min: ICC=0.37, –0.02). During MOD, the 30-s window remained unreliable (ICC=0.32, –0.01), while the 1-min window achieved moderate reliability (ICC=0.63, 0.00). During TT_30_, both shortened windows substantially improved performance (30s: ICC=0.78, –0.02; 1min: ICC=0.95, –0.01), with the 1-min window achieving excellent reliability.

**Conclusion:** DFAa1 analysis window validity is intensity-dependent, with shortened windows showing progressively improved agreement as exercise intensity and heart rate increases. While the 2-min window remains essential for low-intensity monitoring, 1-min or 30-s windows provide appropriate validity during high-intensity exercise, enabling more-responsive real-time feedback. These results support adaptive windowing strategies that dynamically adjust window length based on exercise intensity and the number of included data points, to optimize the analytical validity-temporal responsiveness trade-off.

## Introduction

The application of detrended fluctuation analysis alpha-1 (DFAa1) in load monitoring has emerged as a promising approach for real-time exercise intensity assessment and training optimization (Rogers & Gronwald, 2022). DFAa1, a non-linear index of heart rate (HR) variability (HRV), has been proposed as a biomarker for acute exercise intensity distribution, addressing limitations inherent in traditional methods for determining intensity thresholds (Gronwald et al., 2020). The theoretical foundation of DFAa1 lies in its ability to quantify the fractal-like correlation properties of HR fluctuations, which systematically change with exercise intensity. During low-intensity exercise, DFAa1 values typically remain above 0.75, while increasing exercise intensity causes DFAa1 to decline below this threshold. This behaviour has led to DFAa1 being proposed as a reliable estimate of physiological transitions with several studies reporting strong associations with traditional ventilatory and lactate thresholds (Cassirame et al., 2025; Schaffarczyk et al., 2023; Sempere-Ruiz et al., 2024; Thiart et al., 2023). However, some other studies report more inconsistent associations, including potential methodological reasons in the implementation of the studies (Hoos & Gronwald, 2025).

Besides the scientific rigor, the practical implementation of DFAa1 monitoring faces a fundamental methodological challenge: balancing analytical validity with temporal responsiveness for real-time feedback applications. The 2-min analysis window has been established as the gold standard for DFAa1 calculations, providing sufficient data points to ensure robust and reliable estimates across various exercise intensities. Especially the length of included data sets is a very important issue concerning signal processing (Eke et al., 2002). The justification for this window length lies in the mathematical approximation used to determine the RR intervals required for a valid analysis; i.e., n = N/10, where n represents the window width and N the total number of data points (Chen et al., 2002; Peng et al., 1995). Accordingly, a window width of n = 4-16 requires approximately 160 RR intervals (data points), which corresponds to roughly 2min of cardiac data at a HR of 80 bpm. Notably, no significant differences were observed when DFAa1 was analyzed using 3-min windows or 200 RR intervals across different intensity levels nor when measured from the beginning or end of an exercise bout (Hautala et al., 2003). This demonstrates that while dataset length is critical in signal processing, small variations at specific exercise intensities do not significantly compromise analytical integrity (Hautala et al., 2003). Yet, the 2-min window presents a significant limitation for dynamic training environments: its inherently low temporal resolution makes it unsuitable for detecting rapid physiological changes or providing timely feedback during high-intensity interval training, fatigue onset, or other contexts requiring immediate responsiveness to systemic changes in physiological demands.

This temporal limitation poses a substantial challenge for practical sports applications. During high-intensity interval training, for example, physiological transitions occur rapidly, and coaches require immediate feedback to optimize training stimulus and prevent excessive physiological responses and fatigue. The 2-min window, while highly reliable, cannot provide the responsiveness necessary for real-time training adjustments in these dynamic contexts. Conversely, shorter analysis windows offer enhanced temporal sensitivity but may compromise validity, particularly during low-intensity exercise where signal variability is greatest (Tanoue et al., 2023). Recent research has highlighted the intensity-dependent nature of DFAa1 validity across different window lengths, with evidence suggesting that the validity of ultra-short-term HRV measures is strongly influenced by both exercise intensity and segment length, with shorter windows becoming increasingly unreliable at moderate and low exercise intensities (Sempere-Ruiz et al., 2024; Tanoue et al., 2023).

This intensity dependency creates a complex optimization problem: the analysis window must be long enough to ensure reliable estimates while remaining short enough to detect meaningful physiological transitions in real-time. Understanding how temporal resolution affects the balance between statistical validity and practical responsiveness is crucial for optimizing DFAa1-based monitoring systems for diverse training applications. The present study addresses this fundamental trade-off by systematically investigating the validity and practical utility of different DFAa1 analysis windows (30s, 1min, and 2min) across varying exercise intensities, from steady-state low- and moderate-intensity continuous training to maximal self-paced time trial efforts. By examining how window length affects agreement with the gold-standard 2-min reference across this intensity spectrum, we aim to provide evidence-informed recommendations for selecting appropriate analysis windows in different training contexts and identify potential strategies for adaptive windowing approaches that optimize both validity and temporal sensitivity.

## Methods

### Ethical Approval

This methodological analysis combined two experiments which were approved by the Ethics Committee of UZ Leuven (S69345 and S6650) and for which all tests were conducted at the Bakala Athletic Performance Center (Leuven, Belgium) and Exercise Physiology Research Group. All participants provided written informed consent prior to their involvement in the study and both studies adhered to the principles outlined in the latest revision of the declaration of Helsinki.

### Participants

The present study incorporated two samples of physically active male participants, both employing identical eligibility criteria. The inclusion criteria required participants to engage in at least five hours of sports per week and to be free from severe cardiac arrhythmias, musculoskeletal injuries, or recent major illnesses. Standardized pre-test conditions were implemented, requiring participants to maintain regular sleep and dietary habits, avoid intensive exercise for 36h preceding test sessions, and abstain from alcohol (24h), caffeine (6h), and heavy meals (2h) prior to the visit. We recruited 24 and 20 healthy recreationally active male participants in the first and second experiment, respectively. Five participants were excluded from the first and two from the second experiment due to lack of motivation (n=1), injury/illness (n=2), insufficient fitness (n=1) and inability to complete the sessions (n=1) and for reasons unrelated to the experimental procedures (n=2). Consequently, 19 participants were analysed in the first experiment (mean ± SD: age 22.6 ± 3.6years, height 1.82 ± 0.07m, body mass 71.6 ± 7.6kg, maximal oxygen uptake [VO_2max_] 56.2 ± 6.3ml/kg/min) and 18 participants in the second experiment (mean ± SD: age 21.3 ± 2.6years, height 1.81 ± 0.04m, body mass 73.7 ± 8.5kg, VO_2max_ 55.7 ± 5.7ml/kg/min).

Sample size estimation was performed using G*Power (version 3.1.9.6), based on previously reported effect sizes and correlation values (Fleitas-Paniagua et al., 2024; Kaufmann et al., 2023; Schaffarczyk et al., 2023). Required sample sizes ranged from 12 to 21 (α = 0.05, power = 0.8), indicating that the final cohort size remained adequate despite dropouts.

### Experimental Design

The *first experiment* studied the acute physiological responses to two workload-matched continuous cycling-based training protocols: low-intensity training (LOW) and moderate-intensity training (MOD). Training sessions were matched for workload (kJ) by adjusting time via the formula *E*=*P*×*Δt*. Each participant completed both sessions in a randomized order, with each session separated by a minimum recovery period of seven days to avoid carryover effects (Fig. 1).

**Figure 1:**
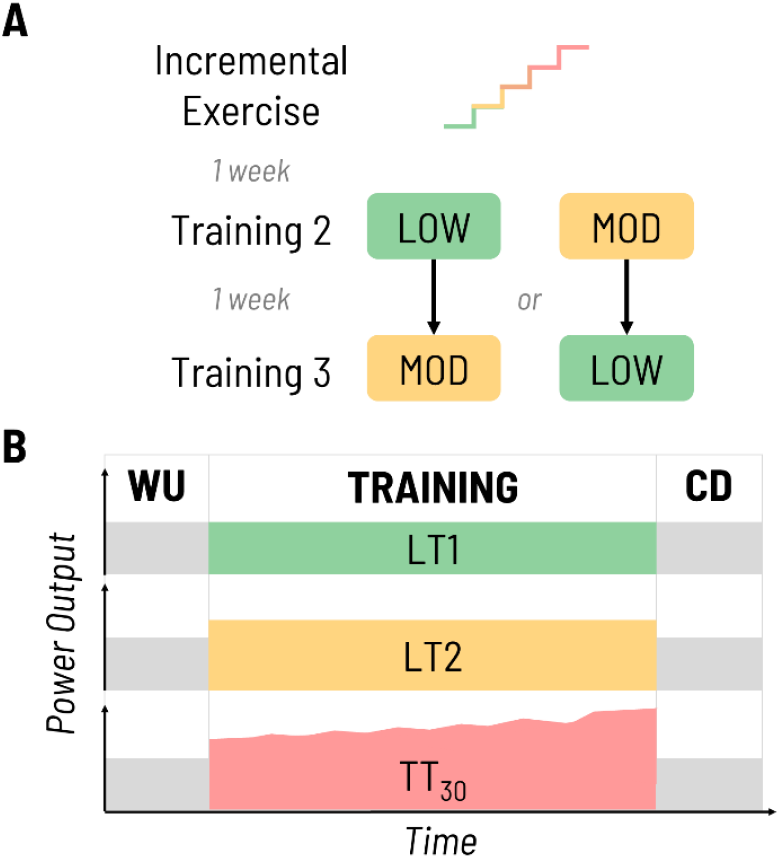
Schematic representation of the experimental design. A) First experiment – The incremental cycling test consisted of 4-min intervals until voluntary exhaustion. Participants where then randomly assigned to two workload-matched continuous cycling protocols: low-intensity training (LOW) and Moderate-intensity training (MOD), at intensities corresponding to the first and second lactate threshold (LT1 and LT2), respectively, including a warm-up (WU) and cool-down (CD). Each session was separated by a minimum of seven days to avoid carryover effects. B) Second experiment – A 30-min self-paced cycling time trial (TT30) with the intention to achieve the highest possible mean power output over 30min.

Both sessions followed an identical three-phase structure: a standardized 10-min warm-up (WU), a main training block of variable intensity and duration depending on the prescribed protocol, and a standardized 10-min cool-down (CD). The WU and CD were performed at the first lactate threshold (LT1). The main training block was performed at intensities corresponding the first and second lactate threshold (LT1 and LT2) for LOW and MOD, respectively (Fig. 1, Table 1). These lactate thresholds and corresponding intensities were determined for each participant a few days before the start of the training protocols during an incremental exercise test. The incremental test started at 40W and increased by 30W every 4min until voluntary exhaustion. Lactate values obtained during these tests were plotted using a third-degree polynomial regression via exphyslab.com (Keir et al., 2022). LT1 was defined as the point at which blood lactate concentration increased by 0.5 mmol/L above baseline (Jamnick et al., 2018; Keir et al., 2022). LT2 was determined using the Log-Log Modified DMax (Log-Poly-ModDMax) method. This method identifies LT2 as the point on the third-degree polynomial curve with the greatest perpendicular distance to the straight line connecting the maximal lactate value and the log-log LT1 (Jamnick et al., 2018). This method was shown to provide the most accurate estimation of the maximal lactate steady state (MLSS) from lactate values measured during a 4-min protocol in trained cyclists (Jamnick et al., 2018).

**Table 1:**
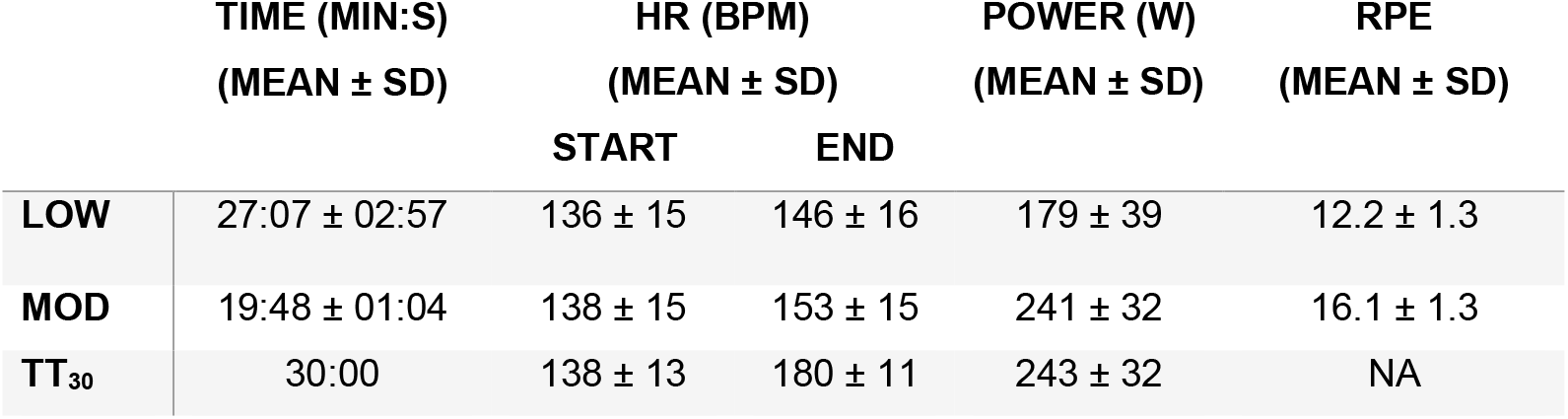
Descriptive overview of the training protocols, detailing the mean duration, heart rate (HR), power output and rating of perceived exertion (RPE) of the low-intensity training (LOW), moderate-intensity training (MOD) and the time trial (TT_30_). HR data are reported separately for the START (first 10min) and END (final 10min) periods of each condition. NA denotes data not available.

|  | TIME (MIN:S)<br>(MEAN ± SD) | HR (BPM)<br>(MEAN ± SD) |  | POWER (W)<br>(MEAN ± SD) | RPE<br>(MEAN ± SD) |
| --- | --- | --- | --- | --- | --- |
|  |  | START | END |  |  |
| **LOW** | 27:07 ± 02:57 | 136 ± 15 | 146 ± 16 | 179 ± 39 | 12.2 ± 1.3 |
| **MOD** | 19:48 ± 01:04 | 138 ± 15 | 153 ± 15 | 241 ± 32 | 16.1 ± 1.3 |
| **TT<sub>30</sub>** | 30:00 | 138 ± 13 | 180 ± 11 | 243 ± 32 | NA |

The *second experiment* originally examined the effect of a nutritional intervention during an eight-week training program and applied a 30-min self-paced cycling time trial (TT_30_) as a baseline performance measure. Participants first completed a ∼10-min WU (75% VO_2max_ + 5min: 85% VO_2max_) followed by a short, non-standardized resting period before the start of the TT_30_. The goal of this test was to achieve the highest possible mean power output over 30min. Every 5min participants were allowed to adjust their pacing strategy and during the final 5min, pacing could be adjusted at one-minute intervals (Poffé et al., 2019) (Fig 1).

For both experiments, these sessions were performed on the same electromagnetically braked cycling ergometer (Cyclus 2, Avantronic, Leipzig, Germany), using either the participant’s own bicycle or a standardized frame. Sessions were scheduled at the same time of day (±2h) to control for circadian influences. Participants remained seated throughout the sessions and no time trial or standing positions were permitted. A fan placed one meter in front of the participant provided consistent airflow across sessions.

## Data Collection

### Heart Rate and RR Interval Measurement

HR and RR interval time series were continuously recorded during the incremental exercise test and training sessions with a Polar H10 chest strap attached sensor (Polar Electro Oy, Kempele, FI, sampling rate: 1000Hz) (Schaffarczyk et al., 2022). The HR sensor was bluetooth-paired with a Polar Pacer watch, which was subsequently connected to Polar Flow for offline analysis. The placement of the HR sensor was standardized for each participant across study visits and HR belt electrodes were moistened to improve conductivity. To ensure accurate signal detection and adequate DFAa1 values, FatMaxxer (Android available, https://github.com/IanPeake/FatMaxxer) was used to verify signal quality prior to each test. Sufficient signal quality was based on R-peak voltage (>1000uV). One participant required repositioning of the HR sensor (1cm to the left from the central position) to achieve sufficient signal quality. HR belt positioning for all participants was standardized across visits by measuring the distance between the HR strap module and the xiphoid process of the sternum.

### Lactate measurements and Rate of Perceived Exertion

Capillary blood lactate was measured using the Lactate Pro 2 analyzer (Arkray, Amstelveen, NL) from a 0.3µL blood sample collected from the earlobe. Rating of perceived exertion (RPE) was assessed using the Borg scale (6–20) (Borg, 1982). During MOD and LOW sessions, RPE was obtained at 5-minute intervals.

## Data Analysis

Clean and artifact-corrected RR interval data were extracted from CSV files for each participant and protocol (i.e., LOW, MOD, TT_30_). Artifact detection and correction were performed using an adaptive regression-based method with a 30-beat sliding window and a deviation threshold of ±10%, allowing a maximum of 10 consecutive corrections to remove ectopic and erroneous beats. A 15-min segment was selected beginning 5min after the onset of the training block or TT_30_ to allow for stabilization of physiological responses. DFAa1 was subsequently computed over this 15-min steady-state segment for the trainings and TT_30_ using a moving-window approach with a window length of 30s (or 1min/2min) and a step size of 1s. For each window, RR intervals were formatted into interbeat interval (IBI) files and pre-processed using detrending, smoothing, interpolation and mean correction. Nonstationary trends were removed using the smoothness priors detrending method (smoothing parameter = 500; cutoff frequency = 0.035Hz). Additionally smoothing was applied using locally weighted regression (LOESS) with a span of 5 points and linear polynomial detrending was performed. Multiscale denoising was conducted using a Daubechies wavelet decomposition with 6 levels. RR intervals were interpolated using cubic spline interpolation and resampled at 2Hz to obtain evenly spaced time series prior to nonlinear analysis. Windows containing fewer than 10 beats or invalid values were excluded from further analysis. DFAa1 was then established over a range of 4-16 beats. Time-resolved DFAa1 values were extracted for each window and used for subsequent statistical analysis. This method enabled a continuous estimation of DFAa1 over time for each participant.

Three window lengths were evaluated for the calculation of DFAa1: 30s, 1min, and 2min. For each method, mean DFAa1 values were computed across all participants, facilitating direct comparison between the shorter windows and the conventional 2-min reference.

## Statistical Interpretation

To evaluate the agreement and validity between the shorter DFAa1 time windows and the 2-min reference, several statistical measures were used. Intraclass Correlation Coefficient (ICC) was calculated to assess the relative validity between methods, indicating the degree to which individual measurements remained consistent across conditions. The ICC values were interpreted according to the guidelines proposed by Koo and Li (2016): < 0.50 = poor reliability, 0.50–0.75 = moderate reliability, 0.75–0.90 = good reliability, and >0.90 = excellent reliability. A two-way random-effects model with absolute agreement was used for all ICC calculations. To assess absolute agreement, Bland-Altman plots were generated and the mean bias along with 95% limits of agreement (LoA) were reported. The bias represents the average difference between methods, while the LoA (mean difference ± 1.96 × SD) indicate the expected range within which most differences between methods fall. Standard Error of Measurement (SEM) was used to assess absolute validity, reflecting the amount of random error in repeated measurements. It was calculated using the formula: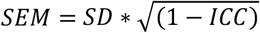, where SD is the standard deviation. Furthermore, the Smallest Worthwhile Change (SWC) was calculated as 0.2 times the pooled standard deviation of the combined measurements, in line with Cohen’s criteria for a small effect size. All statistical analyses were performed using custom-built MATLAB scripts to allow for efficient and standardized batch processing across all participants and conditions.

## Results

When comparing exercise intensity metrics, participants achieved a mean power output of 179W during the LOW condition, with HR increasing from 136 to 146bpm and a mean RPE of 12.2. In contrast, the MOD condition elicited a mean power of 241W, a HR range from 138 to 153bpm, and a higher mean RPE of 16.1. During the TT_30_ trial, participants produced a mean power of 243W, with HR ranging from 138 to 180bpm (see Table 1).

During *LOW*, the comparison between the 30-s and 2-min methods revealed poor agreement of DFAa1 (ICC = 0.02, 95% CI [0.00, 0.09]) with a mean bias of –0.05, LoA (–0.16 to 0.06), a SEM of 0.04 and a SWC of 0.01. In contrast, the 1-min window showed better but still poor agreement (ICC = 0.37, 95% CI [0.31, 0.43]) with low mean bias of –0.02, narrower LoA (–0.10 to 0.06), a lower SEM 0.03 and a lower SWC (0.01) (Fig. 2).

**Figure 2:**
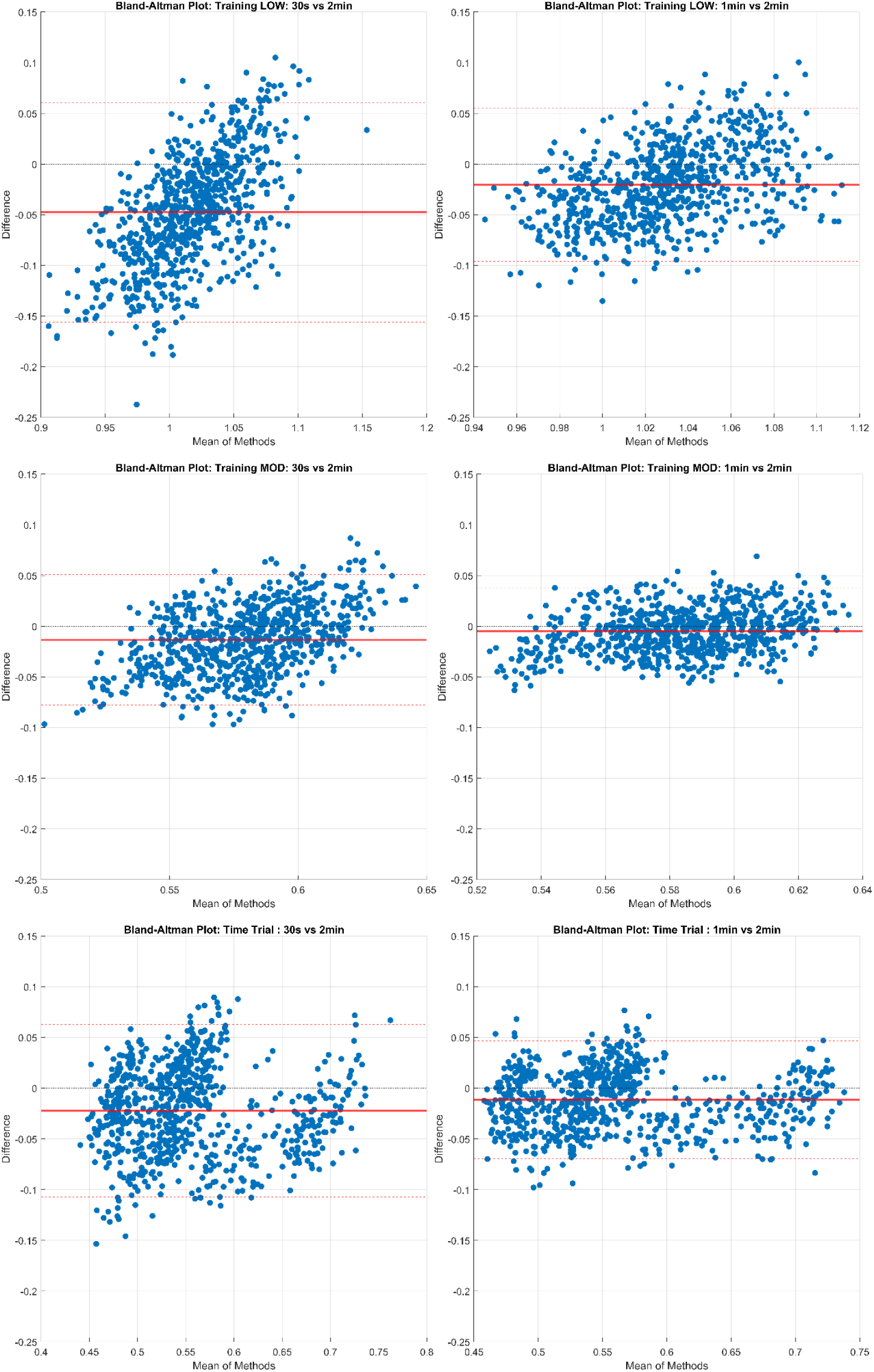
Blant-Altmann plots comparing 30-s (left) and 1-min (right) windows with a 2-min window for the LOW (top), MOD (middle), and Time Trial (bottom) protocols.

For *MOD*, the 30-s window produced poor reliability (ICC = 0.32, 95% CI [0.26, 0.38]) and a smaller mean bias of –0.01, narrower LoA (–0.08 to 0.05), a smaller SEM (0.02) and smaller SWC (0.01) compared to LOW. The 1-min method yielded moderate reliability (ICC = 0.63, 95% CI [0.58, 0.67]) with a negligible mean bias of 0.00, narrower LoA (–0.05 to 0.04), lower SEM (0.02) and SWC (0.01) compared to LOW (Fig. 2).

During the *TT*_*30*_ the 30-s method showed good reliability (ICC = 0.78, 95% CI [0.75, 0.81]) with a mean bias of –0.02 narrow LoA (–0.11 to 0.06), a SEM of 0.03 and a SWC of 0.015. The 1-min window achieved excellent reliability (ICC = 0.95, 95% CI [0.89, 0.92]) with a mean bias of -0.01, narrow LoA (–0.07 to 0.05), a SEM of 0.02 and a SWC of 0.01 (Fig. 2).

## Discussion

The current findings reveal a fundamental methodological challenge in DFAa1 implementation: the inherent trade-off between analytical validity and temporal responsiveness. While the 2-min window is currently the most commonly used approach, its low temporal resolution renders it unsuitable for applications requiring rapid detection of physiological changes, such as response and fatigue monitoring during high-intensity interval training or real-time intensity adjustments. This investigation systematically quantified the intensity-dependent nature of window validity and identified conditions under which shorter analysis windows may provide acceptable alternatives to the gold-standard 2-min reference.

### Intensity-Dependent Window Performance

The intensity-dependent nature of window validity represents a key finding of this investigation. *During low-intensity exercise* (LOW protocol at LT1), both 30-s and 1-min windows demonstrated poor agreement with the 2-min reference, with the 30-s method showing extremely poor reliability (ICC = 0.02, 95% CI [0.00, 0.09]) and the highest SEM observed across all conditions (0.04). While this SEM represents approximately 4.9% of the typical DFAa1 range (0.3–1.3) and might be considered technically acceptable in absolute terms, the combination of poor relative agreement (ICC) and wide LoA (–0.16 to 0.06) renders the 30-s window unsuitable for low-intensity contexts. The 1-min window showed improved but still poor agreement (ICC = 0.37, 95% CI [0.31, 0.43]), with narrower LoA (–0.10 to 0.06) and lower SEM (0.03), suggesting it represents a more viable alternative than the 30-s window, though neither can be recommended as reliable substitutes for the 2-min reference during low-intensity exercise.

*During moderate-intensity exercise* (MOD protocol at LT2), shorter window performance improved substantially, though clear differences remained between the 30-s and 1-min methods. The 30-s window continued to show poor relative reliability (ICC = 0.32, 95% CI [0.26, 0.38]), albeit with a smaller mean bias (–0.01), narrower LoA (–0.08 to 0.05), and smaller SEM (0.01) compared to LOW conditions. The 1-min window achieved moderate reliability (ICC = 0.63, 95% CI [0.58, 0.67]) with negligible bias (–0.00), narrow LoA (–0.05 to 0.04), and the lowest SEM observed across all conditions (0.02). These results suggest that at moderate intensities corresponding to the second lactate threshold, the 1-min window begins to provide acceptable agreement with the 2-min reference, though the 30-s window remains insufficiently reliable for practical application.

The most striking improvement in shorter window performance occurred *during the 30-min maximal time trial* (TT_30_), in which HR progressively increased toward exhaustion. Under these high-intensity conditions, both shorter windows demonstrated substantially improved agreement with the 2-min reference. The 30-s window achieved good reliability (ICC = 0.78, 95% CI [0.75, 0.81]) with narrow LoA (–0.11 to 0.06) and SEM of 0.03, while the 1-min window demonstrated excellent reliability (ICC = 0.95, 95% CI [0.89, 0.92]) with even narrower LoA (–0.07 to 0.05) and SEM of 0.02. This marked improvement likely reflects the reduced variability in DFAa1 calculations as exercise intensity increases and HR becomes more constrained by physiological limitations.

### Physiological Mechanisms Underlying Intensity-Dependent Validity

The intensity-dependent improvement in shorter window validity can be understood through the physiological mechanisms governing HR dynamics across the exercise intensity spectrum. At low intensities, cardiovascular regulation involves complex interactions between parasympathetic and sympathetic nervous system activity, baroreceptor reflexes, respiratory sinus arrhythmia, and various other regulatory mechanisms that produce intricate, long-range correlations in the HR signal (Perini & Veicsteinas, 2003). These multifaceted regulatory inputs create a highly variable signal that requires sufficient data points (i.e., extended observation periods) to accurately characterize these fractal dynamics. Shorter windows simply do not capture enough cardiac cycles to reliably estimate these correlation properties, resulting in substantial random variation around the true DFAa1 value (Michael et al., 2017). As exercise intensity increases toward and beyond LT2, parasympathetic withdrawal increases, sympathetic drive dominates, and the regulatory complexity diminishes (Rogers & Gronwald, 2022). The HR signal becomes increasingly constrained by maximal cardiovascular output limitations, reducing the variability and fractal complexity that necessitate longer analysis windows (Shaffer et al., 2020; Tanoue et al., 2023).

This physiological transition explains why the 30-s window, which showed extremely poor reliability during LOW (ICC = 0.02), achieved good reliability during TT_30_ (ICC = 0.78) - a nearly 40-fold improvement in relative agreement. Similarly, the 1-min window progressed from poor reliability during LOW (ICC = 0.37) to excellent reliability during TT_30_ (ICC = 0.95). These findings align with recent research demonstrating that ultra-short-term HRV measures function as “proxies of proxies” whose validity depends critically on the underlying physiological state and signal characteristics (Shaffer et al., 2020; Tanoue et al., 2023).

### Practical Implications and Future Directions

These findings demonstrate that no single fixed analysis window is universally optimal across exercise contexts. The 2-min window, while providing superior validity across all intensities, cannot provide timely feedback during dynamic training scenarios such as high-intensity interval training, where a 2-min delay in detecting DFAa1 changes could result in excessive physiological demand and fatigue accumulation. Conversely, shorter windows show insufficient validity during low-intensity training - which constitutes the majority of training volume in endurance programs - risking misinterpretation of random signal variation as meaningful physiological changes.

The substantial improvement in shorter window validity at high intensities suggests that context-dependent windowing strategies may offer practical solutions. Adaptive algorithms could dynamically adjust window length based on availability of datapoints (i.e., number of IBI), for instance employing 2-min windows with lower HR and transitioning to shorter windows (e.g. 1-min or 30-s windows) with higher HR – where ∼140-145bpm could serve as a provisional threshold. Such internal load-based switching would preserve validity during low-intensity training while enabling rapid detection of intensity transitions during higher-intensity efforts. However, implementing threshold-based adaptive strategies requires careful validation to ensure benefits outweigh potential artifacts from dynamic window adjustment.

### Methodological Considerations

Several limitations warrant discussion. *First*, the moving window approach with 1-s increments introduces temporal overlap between successive calculations, potentially inflating apparent agreement between methods. Future research should examine whether non-overlapping window approaches yield different validity estimates, particularly during intensity transitions.

*Second*, this study focused exclusively on cycling exercise in controlled laboratory conditions with recreationally active males (VO_2max_ ∼56 mL/kg/min). Generalizability to other exercise modalities (running, rowing, swimming), field-based environments, female athletes, higher-level athletes, or clinical populations requires verification. Different exercise modes may introduce distinct movement artifacts or respiratory patterns, and physiological coupling mechanisms affecting DFAa1 values, signal quality and optimal window selection.

*Third*, the study examined three specific protocols (continuous low-intensity, continuous moderate-intensity, and progressive time trial). High-intensity interval training with repeated work-rest transitions, variable-intensity (stochastic) fartlek training, or ultra-endurance efforts may present unique challenges not addressed in this investigation.

## Conclusions

This investigation demonstrates that DFAa1 window validity is strongly intensity-dependent. During low-intensity exercise, neither 30-s nor 1-min windows provide acceptable validity, necessitating 2-min analysis. During moderate-intensity exercise, 1-min windows achieve moderate validity. During self-paced time trail performance up to high exercise intensity, both 1-min (excellent) and 30-s (good) windows demonstrate acceptable agreement with the 2-min reference. Adaptive windowing strategies that adjust analysis window lengths based on exercise intensity capturing the number of included data points depending on IBI represent a promising direction for enhancing practical DFAa1 monitoring across the full intensity spectrum of endurance training.

## Abbreviations

DFAa1: Detrended Fluctuation Analysis alpha-1
HR: Heart Rate
HRV: Heart Rate Variability
VO_2max_: Maximal oxygen consumption
LOW: Low-intensity training
MOD: Moderate-intensity training
WU: Warm-up
CD: Cool-down
LT1: First lactate threshold
LT2: Second lactate threshold
MLSS: Maximal lactate steady state
TT_30_: 30-min self-paced cycling time trial
IBI: Interbeat interval
ICC: Intraclass correlation coefficient
LoA: Limits of agreement
SD: Standard deviation
SEM: Standard error of measurement
SWC: Smallest worthwhile change
RPE: Rating of perceived exertion

## Acknowledgements

We would like to thank the Exercise Physiology Group (KU Leuven) for their support in the data collection process. We would also like to thank Dante Mantini, Yoram Müller-Jabusch, Jef Leplae, Casper Vananderoye and Christophe Dausin for their contributions and insightful discussions.

## Author Contribution Statement

JDM, AO, TG and TdB conceived and designed research. JDM and AO conducted experiments. JDM programmed analytical tools. JDM, AO and TdB analyzed data. JDM, AO, TG and TdB wrote the manuscript. All authors read and approved the manuscript.

## References

Borg, G. A. V. (1982). Psychophysical bases of Perceived Exertion. Medicine and Science in Sports and Exercise, 14(5), 377–381.

Cassirame, J., Eustache, E., Garbellotto, L., Chevrolat, S., Gimenez, P., & Leprêtre, P. M. (2025). Detrended fluctuation analysis to determine physiologic thresholds, investigation and evidence from incremental cycling test. European Journal of Applied Physiology, 125(2), 523–533. 10.1007/s00421-024-05614-z

Chen, Z., Ivanov, P. C., Hu, K., & Stanley, H. E. (2002). Effect of nonstationarities on detrended fluctuation analysis. Physical Review E - Statistical Physics, Plasmas, Fluids, and Related Interdisciplinary Topics, 65(4), 15. 10.1103/PhysRevE.65.041107

Eke, A., Herman, P., Kocsis, L., & Kozak, L. R. (2002). Fractal characterization of complexity in temporal physiological signals. In Physiol. Meas (Vol. 23). 10.1088/0967-3334/23/1/201

Fleitas-Paniagua, P. R., Marinari, G., Rasica, L., Rogers, B., & Murias, J. M. (2024). Heart Rate Variability Thresholds: Agreement with Established Approaches and Reproducibility in Trained Females and Males. Medicine and Science in Sports and Exercise, 56(7), 1317–1327. 10.1249/MSS.0000000000003412

Gronwald, T., Rogers, B., & Hoos, O. (2020). Fractal Correlation Properties of Heart Rate Variability: A New Biomarker for Intensity Distribution in Endurance Exercise and Training Prescription? Frontiers in Physiology, 11. 10.3389/fphys.2020.550572

Hautala, A. J., Mäkikallio, T. H., Seppänen, T., Huikuri, H. V., & Tulppo, M. P. (2003). Short-term correlation properties of R-R interval dynamics at different exercise intensity levels. Clinical Physiology and Functional Imaging, 23(4), 215–223. 10.1046/j.1475-097X.2003.00499.x

Hoos, O., & Gronwald, T. (2025). Detrended fluctuation analysis of heart rate variability during exercise: Time to reconsider the theoretical and methodological background. Comment on: Cassirame et al.s (2025) Detrended fluctuation analysis to determine physiologic thresholds, investigation and evidence from incremental cycling test. Eur J Appl Physiol 125:523–533. European Journal of Applied Physiology. 10.1007/s00421-025-05859-2

Jamnick, N. A., Botella, J., Pyne, D. B., & Bishop, D. J. (2018). Manipulating graded exercise test variables affects the validity of the lactate threshold and V_ O2peak. PLoS ONE, 13(7). 10.1371/journal.pone.0199794

Kaufmann, S., Gronwald, T., Herold, F., & Hoos, O. (2023). Heart Rate Variability-Derived Thresholds for Exercise Intensity Prescription in Endurance Sports: A Systematic Review of Interrelations and Agreement with Different Ventilatory and Blood Lactate Thresholds. In Sports Medicine - Open (Vol. 9, Number 1). Springer Science and Business Media Deutschland GmbH. 10.1186/s40798-023-00607-2

Keir, D. A., Iannetta, D., Mattioni Maturana, F., Kowalchuk, J. M., & Murias, J. M. (2022). Identification of Non-Invasive Exercise Thresholds: Methods, Strategies, and an Online App. In Sports Medicine (Vol. 52, Number 2, pp. 237–255). Springer Science and Business Media Deutschland GmbH. 10.1007/s40279-021-01581-z

Michael, S., Graham, K. S., & Oam, G. M. D. (2017). Cardiac autonomic responses during exercise and post-exercise recovery using heart rate variability and systolic time intervals-a review. In Frontiers in Physiology (Vol. 8, Number MAY). Frontiers Media S.A. 10.3389/fphys.2017.00301

Peng, C.-K., Havlin, -∼ S, Hausdorff, J.-$ J. M., Mietus, § J E, Stanley, H.E., & Goldberger, A. L. (1995). Fractal Mechanisms and Heart Rate Dynamics Long-range Correlations and Their Breakdown With Disease. In Journal of Electrocardiology (Vol. 28).

Perini, R., & Veicsteinas, A. (2003). Heart rate variability and autonomic activity at rest and during exercise in various physiological conditions. In European Journal of Applied Physiology (Vol. 90, Numbers 3–4, pp. 317–325). 10.1007/s00421-003-0953-9

Poffé, C., Ramaekers, M., Van Thienen, R., & Hespel, P. (2019). Ketone ester supplementation blunts overreaching symptoms during endurance training overload. Journal of Physiology, 597(12), 3009–3027. 10.1113/JP277831

Rogers, B., & Gronwald, T. (2022). Fractal Correlation Properties of Heart Rate Variability as a Biomarker for Intensity Distribution and Training Prescription in Endurance Exercise: An Update. In Frontiers in Physiology (Vol. 13). Frontiers Media S.A. 10.3389/fphys.2022.879071

Schaffarczyk, M., Rogers, B., Reer, R., & Gronwald, T. (2022). Validity of the Polar H10 Sensor for Heart Rate Variability Analysis during Resting State and Incremental Exercise in Recreational Men and Women. Sensors, 22(17). 10.3390/s22176536

Schaffarczyk, M., Rogers, B., Reer, R., & Gronwald, T. (2023). Validation of a non-linear index of heart rate variability to determine aerobic and anaerobic thresholds during incremental cycling exercise in women. European Journal of Applied Physiology, 123(2), 299–309. 10.1007/s00421-022-05050-x

Sempere-Ruiz, N., Sarabia, J. M., Baladzhaeva, S., & Moya-Ramón, M. (2024). Reliability and validity of a non-linear index of heart rate variability to determine intensity thresholds. Frontiers in Physiology, 15. 10.3389/fphys.2024.1329360

Shaffer, F., Meehan, Z. M., & Zerr, C. L. (2020). A Critical Review of Ultra-Short-Term Heart Rate Variability Norms Research. In Frontiers in Neuroscience (Vol. 14). Frontiers Media S.A. 10.3389/fnins.2020.594880

Tanoue, Y., Nakashima, S., Komatsu, T., Kosugi, M., Kawakami, Saki, Kawakami, Shotaro, Michishita, R., Higaki, Y., & Uehara, Y. (2023). The Validity of Ultra-Short-Term Heart Rate Variability during Cycling Exercise. Sensors, 23(6). 10.3390/s23063325

Thiart, N., Coetzee, B., & Bisschoff, C. (2023). Heart Rate Variability-Established Thresholds to Determine the Ventilatory and Lactate Thresholds of Endurance Athletes. International Journal of Human Movement and Sports Sciences, 11(2), 398–410. 10.13189/saj.2023.110217

